# Genotype-dependent behavioral signatures of opioid withdrawal revealed by automated behavioral quantification

**DOI:** 10.64898/2026.10.01.756016

**Authors:** Jacob A. Beierle, Gautam Sabnis, Vivek Kumar

**Author notes:** Senior author (Vivek Kumar).

## Abstract

Opioid withdrawal drives continued opioid use in opioid use disorder (OUD), which affected an estimated 4.8 million Americans in 2024. However, only one non-opioid medication is FDA-approved for withdrawal treatment, and there is great need for more diverse treatment options. Mice are powerful models to study opioid withdrawal because they allow for controlled drug and environmental exposure, express OUD-relevant phenotypes, and provide access to withdrawal-relevant tissue at exact time points. However, mouse models are hindered by their over reliance on a single genetic background (C57BL/6J). Using a single background risks limiting the translatability of preclinical studies as they are projected into genetically diverse human populations. Inferring withdrawal intensity in mice typically relies on single behaviors or indexes of behaviors, and applying these approaches across mouse genetic diversity faces a critical challenge: differing withdrawal responses could either encode differences in withdrawal severity, or differences in withdrawal-evoked behaviors independent of severity. To overcome this limitation, a more comprehensive assessment of opioid withdrawal behaviors across mouse genetic diversity is needed. Here we leverage computer vision and machine learning to automate the simultaneous quantification of many mouse behaviors during opioid withdrawal in eight diverse strains of mice. We reveal that all eight strains exhibit withdrawal-evoked behaviors, and these behaviors are sufficient to identify the withdrawal state of the mouse. However, the effect of withdrawal on individual behaviors or indexes of behaviors is not uniform across genetic diversity, and extending methods tuned in one strain risks misclassifying withdrawal in others. Finally our approach was able to distinguish graded withdrawal intensity, as defined by opioid exposure within 3 of 4 strains tested. We therefore conclude that opioid withdrawal behavior is profoundly shaped by genetic background, but withdrawal severity can nonetheless be established within a background. Combining automated computer-vision phenotyping with genotype-aware modeling provides a scalable path toward genetically informed withdrawal assessment in diverse mouse populations, with the ultimate goal of identifying the genetic and biological networks that modulate opioid withdrawal and may represent new targets for treating opioid use disorder.

## 1 Introduction

### 1.1 Improving withdrawal treatment is a pathway to better OUD outcomes

In 2024, an estimated 4 million Americans suffered from opioid use disorder (OUD) [48], a psychiatric disorder characterized by compulsive chronic use of opioids despite negative repercussions [1]. Only *∼*17% of people with OUD received medication for opioid use disorder (2017) [21, 48], a treatment gap that contributed to the nearly 55,000 opioid overdose deaths in 2024 [32]. Opioid withdrawal, the negative affective and somatic consequences of opioid abstinence, represents a clinically important opportunity to intervene during OUD. Withdrawal severity varies substantially between patients [47], increases treatment dropout [47] and relapse [33, 42], and treating opioid withdrawal improves retention in OUD treatment [14]. There is only one FDA-approved non-opioid treatment for withdrawal despite its varied neuroanatomical [6, 15, 23, 26, 27, 29, 44, 46, 52, 55] and mechanistic [9, 12, 28, 37, 38, 43, 49] effectors, implying broader opportunities for intervention. Diversifying withdrawal treatment, therefore, represents an opportunity to improve OUD patient outcomes.

Animal models are needed to better understand opioid withdrawal because they allow for controlled drug exposure and environment as well as access to withdrawal relevant tissue. Specifically, mice are a valuable model system for studying OUD because of their shared OUD behaviors like escalating opioid consumption [50], tolerance [53], and withdrawal [25], alongside substantial but controlled genetic diversity [57]. In rodents, withdrawal is studied either by cessation of opioid exposure (spontaneous withdrawal), or through pharmacological precipitation (naloxone-precipitated withdrawal [53]) and the severity of opioid withdrawal scales with the magnitude of prior opioid exposure [10, 35, 36]. Withdrawal intensity can be inferred through individual behaviors or behavioral indices (e.g. jumping, tremor, rearing), physiological markers, anxiety measures, hyperalgesia, and reward assays. Together, these strengths have empowered mouse models to uncover distinct neuroanatomy, cellular processes, and neurotransmitter systems regulating opioid withdrawal.

Despite the power of mice as models of opioid withdrawal, they are limited by an over-reliance on a single genetic background (C57BL/6J) in two primary ways. First, using one isogenic background to study the genetics, neuroscience, and pharmacology of opioid withdrawal risks limiting the translatability of preclinical studies as they are projected into genetically diverse human populations. Expanding withdrawal research into genetically diverse mice would increase confidence in our preclinical findings, but few measures have been developed and validated for diverse genetic backgrounds. It is therefore unclear what behaviors discriminate withdrawal severity across diverse genetic backgrounds. Second, OUD phenotypes are influenced by genetics in mice [2,5,13], like they are in humans [4, 8, 18, 51, 58], and understanding the drivers of that genetic risk grants insight into the biological systems driving withdrawal. Accessing the naturally occurring genetic modulators of opioid withdrawal is not possible using one isogenic strain.

The influence of genetic background on opioid withdrawal has been studied in mice [19, 20], but has largely relied on the single phenotype of jumping, which is not genetically correlated with other opioid withdrawal outcomes like hypothermia or fecal boli production [30]. This raises the question, are these studies interrogating genetic predisposition to opioid withdrawal severity, or the genetic propensity to jump during opioid withdrawal? Addressing these limitations requires a more multivariate understanding of the behavioral response to opioid withdrawal across genetics from which we can build genotype-aware models.

Accessing this more multivariate behavioral response presents a serious technical challenge. Manual scoring is laborious and time consuming, limiting the number of behaviors, the number of animals, and duration of monitoring available to a study. Additionally, commonly used composite withdrawal scores have been developed and weighted in a small number of genetic backgrounds, most often C57BL/6J, and like single measures, it is unclear how representative they are of withdrawal across genetic diversity. Higher-throughput alternatives like videography-based centroid tracking trade dimensionality for scale and thereby lose the ability to score a large range of complex behaviors. Additionally, many assays indexing the affective and physiological dimensions of withdrawal yield a single measurement per animal and cannot be acquired concurrently, similarly limiting the dimensionality accessible during phenotyping.

Computer vision and machine learning address these constraints in several ways. Neural-network based segmentation and pose estimation automate the complex representation of an animal, and using the spatial and temporal relation of these data allows access to features not feasibly scorable by hand at scale, like gait and body geometry. These keypoints can also be leveraged for supervised behavior classification, eliminating inter-rater variability by encoding and reusing expert annotations [7]. Because of the low cost to predict these behavioral features it is feasible to quantify non-canonical behaviors alongside canonical ones, allowing data, not convention, to indicate which features separate withdrawal states in each background. Together this allows scalable access to rich behavioral indices with high temporal and spatial resolution, making it feasible to measure the high-dimensional behavior space needed to build genotype-informed models of withdrawal. Computer vision and machine learning behavioral indices have been leveraged to quantify frailty/biological age [16, 40], seizure [41], pain [3, 34], and social defeat [24], highlighting their value in behavioral neuroscience.

In this work we characterize the behavioral expression of precipitated opioid withdrawal across eight genetically diverse inbred strains of mice, selected for their historic representation in the withdrawal field and inclusion in genetically diverse panels of mice. We employ naloxone precipitated withdrawal for the precise induction of withdrawal and to overcome the unknown effect of genetics on the kinetics of spontaneous withdrawal. To access high-dimensional behavior data we leverage the JAX Animal Behavior System (JABS) [7], an integrated computer vision/machine learning phenotyping platform that streamlines behavior testing, video collection, and behavior analysis. Through this approach we observe genotype distinct patterns of withdrawal-associated behavior and discriminate the withdrawal condition in all eight strains of mice. We also demonstrate the ability of high-dimensional behavioral indices to resolve graded withdrawal responses within genetic backgrounds, defined through escalating opioid exposure paradigms. Finally, in both approaches, we observe a core set of features that predicts withdrawal condition across genetics, suggesting a partially conserved behavioral response to opioid withdrawal. Our results provide the groundwork for high-dimensional, genetics-informed behavioral models of opioid withdrawal that can serve as the basis for investigations into genetic, pharmacological, and neural effectors of opioid withdrawal.

## 2 Methods

### 2.1 Mice

Mice were obtained from The Jackson Laboratory (Bar Harbor, ME), at 10–14 weeks of age and included 129S1/SvImJ (#002448), A/J (#000646), AKR/J (#000648), BALB/cJ (#000651), C57BL/6J (#000664), CAST/EiJ (#000928), DBA/2J (#000671), NOD/ShiLtJ (#001976). Mice were housed in a 12 hr light/dark vivarium, provided ad lib 5K52 Lab Diet sterilized grain and acidified water, and were acclimated to the vivarium for at least 7 days before testing. All animal protocols were approved by and adhered to JAX IACUC guidelines. In total, 142 mice were included in this study. Sample sizes by strain, sex, and treatment can be found in supplemental table S1, S2.

### 2.2 Induction of dependence and withdrawal testing

Mice fell into 3 groups: baseline mice, withdrawal mice, and high withdrawal mice, with some mice tested in both baseline and withdrawal conditions. Withdrawal mice were given once daily injections of morphine sulfate (Cayman Chemical, #25955; i.p., 10mL/kg body weight) 2 hours after lights on (08:00) in sterile saline for 5 consecutive days. Each day the dose was increased 10 mg/kg, starting at 10mg/kg on day 1, and ending at 50 mg/kg on day 5. On day 5, 2 hours following the final injection of morphine, mice were injected with 10 mg/kg naloxone (Millipore Sigma, #N7758; i.p., 10mL/kg body weight), to precipitate withdrawal. High withdrawal mice received morphine twice daily (at 08:00, 16:00) for 4 days, starting with 20 mg/kg and escalating 20 mg/kg/day to 80 mg/kg. On day 5, high withdrawal mice received a single injection of 100 mg/kg morphine (08:00), followed 2 hours later by 10 mg/kg naloxone. Baseline mice received 2 injections of saline (10ml/kg body weight) on day 5, separated by 2 hours.

Behavioral testing was conducted immediately following the second injection on day 5 (saline or naloxone). Mice were habituated to the testing room 30 mins prior to testing, then placed in 20.5 in x 20.5 in open field arenas as previously described [11], and 150 min of top-down videography was captured using the JABS system [7].

Mice that received both treatments were tested for baseline behaviors 3-4 days before morphine administration, then administered morphine and tested as described.

### 2.3 Video processing and behavioral predictions

The video capture methods, tracking, and segmentation networks have been described previously [11, 22]. In brief, a neural network trained to generate a mouse segmentation mask produced an ellipse fit in each frame, along with a track. Additionally, 12 keypoints were estimated in each frame using a deep convolutional neural network [45]. The keypoints captured include the nose, left ear, right ear, base of the neck, left forepaw, right forepaw, mid-spine, left rear paw, right rear paw, base of the tail, mid-tail, and tip of the tail. For each key point at each frame, an x-coordinate, y-coordinate, and confidence score were recorded. A minimum confidence threshold of 0.3 was applied to determine which keypoints were included in subsequent analyses.

Ellipse fit tracking was used to infer locomotion and traditional open field measures as previously described [11]. Key point predictions were used to infer gait and spinal posture (morphometric) features [16, 45]. Keypoint and segmentation data were used to train supervised classifiers using the JABS python-based GUI [7]. Jumping and freezing classifiers were trained specifically for these experiments, validated through k-fold cross validation, and their accuracy ( (true positives + true negatives) / all observations) and F1 scores (harmonic mean of precision and recall) are reported in Table S3. A complete list of all calculated features, and their definitions, can be found in supplemental materials.

### 2.4 Feature processing and imputation

Prior to each dimensionality reduction analysis, behavioral feature data were subjected to a series of qualitycontrol and preprocessing steps. Features missing more than 25% of observations or with fewer than 3 observations within any strain-by-treatment group were excluded from analysis. All other missing features were imputed based on the median of the strain*tx observations. To calculate delta features (withdrawal baseline), mice missing one observation were completed using median imputation of that strain*treatment group. After imputation, zero-variance features and features with one differing observation were removed from the feature matrix to prevent instability in subsequent dimensionality reduction analysis and k-fold cross validation.

When producing subsequent models, the remaining numeric features within the subset of mice in each model were z-score transformed (centered and scaled to unit variance) column-wise across all observations. The zscored feature matrix served as the common input for sPCA and sPLS-DA methods. These models included an “all mice” model, a “high activity” mouse model (including C57BL/6J, CAST/EiJ, DBA/2J, and NOD/ShiLtJ strains), a “low activity” strain model (including 129S1/SvImJ, A/J, AKR/J, and BALB/cJ strains), and 8 strain specific models.

### 2.5 Sparse principal component modeling

Sparse PCA (sPCA) was used to summarize the curated behavioral feature set in an unsupervised, low-dimensional space. Models were fitted with mixOmics [39] to thirteen feature matrices: per-mouse withdrawal-minus-baseline deltas, all sessions pooled, high- and low-activity strain groups, each of the eight strains separately, and baseline sessions only. First, a search ceiling K_max was set as the smallest number of dense principal components reaching 77.5% cumulative variance in the same z-scored matrix (capped at min(n, p) − 1). For each component, the number of features per component (keepX) was selected by mixOmics::tune.spca using 5-fold cross-validation with 5 repeats (folds reduced to floor(n/3) for models with fewer than 15 observations) from a candidate vector of proportions of each model’s own feature count (0.025, 0.05, 0.1, 0.2, 0.4, 0.6 and 0.8, rounded to integers). Components were then retained only if their cross-validated reproducibility was significantly greater than that model’s own chance level, defined by the average null value over each fold. The mean of the five repeat-level means for each component was compared with this chance level by a one-sided, one-sample t-test on four degrees of freedom, and components were retained up to the first that failed at *α* = 0.10 (matching the *α* mixOmics uses for keepX selection). The sPCA model was fit using these parameters using the Sparse Non-linear Iterative Partial Least Squares algorithm implemented in mixOmics, with a maximum of 1,000 iterations. A scatter plot of PC1 versus PC2 scores was used to quantitatively assess separation of withdrawal or strain groups.

### 2.6 Sparse partial least squares discriminant analysis modeling

sPLS-DA was performed using the mixOmics::splsda function from the mixOmics R package [39] as a supervised complement to the unsupervised sPCA using baseline and withdrawal observations. The model was fit on the same z-scored feature matrix used for sPCA, and outcome vairable defined as treatment group. The number of features per component (keepX) was selected using tune.splsda (5-fold × 5 repeats, centroids.dist, minimising BER) with folds capped at the smallest treatment size, from a candidate vector of proportions of each model’s own feature count (0.025, 0.05, 0.1, 0.2, 0.4, 0.6 and 0.8, rounded to integers). The number of components was selected using the selected ncomp, and this selected number of components was used to assess within and cross model performance. All sPLS-DA results were visualized as scatter plots of discriminant component scores plotted in the component 1 versus component 2 space.

### 2.7 Evaluation of within and cross model performance

To determine the performance of each model, and whether the behavioral signatures learned by one model generalize to other subsets, a systematic within and cross-group prediction analysis was conducted. Each comparison was assessed by subject-grouped, class-stratified k-fold cross-validation, where folds are assigned by mouse (so a mouse’s baseline and withdrawal rows never straddle training and test sets), and are stratified on treatment class. 10 folds were requested, but capped at the number of mice carrying the rarer class. Model performance was assessed as an 11 * 11 matrix asking how well a model trained on one group classifies withdrawal against baseline in another, with the on diagnal being within model performance, and off diagonal cross model performance.

Within model performance was assessed by refitting the sPLS-DA with the model’s tuned keepX and component count. Centering and scaling are re-derived from that fold’s training rows only and held-out observations are classified by centroid distance (‘centroids.dist‘), along those kept components. Predictions were pooled across folds into a single confusion matrix. Accuracy, balanced error rate (BER), sensitivity, specificity, precision and F1 were computed from that pooled matrix with withdrawal as the positive class.

Cross model performance (each of the 110 off-diagonal cells) was assessed by tuning and fitting its own sparse sPLS-DA on the training group’s observations using the methods reported above and was restricted to features that pass the missing-data filter in both the training and the test group and have non-zero variance in the training group. Test-group features are z-scored with the training group’s means and SDs, so no test-set information enters the scaling, and test observations are classified by centroid distance against the training class centroids. Because the eleven groups are nested (the pooled group contains the activity groups, which contain the strains), offdiagonal cells follow 3 strategies to ensure performance is assessed leakage-free. Group 1: The two groups share no mice (e.g. A/J tested on DBA/2J). The model is trained on the full training group and the whole test group is projected into it and classified in one pass. Group 2: The training group’s own mice sit inside the test set (e.g. C57BL/6J tested on All). Those mice are removed from the test set before projection, so a model is never scored on an animal it trained on. Group 3: Test mice are part of the training group (e.g. All tested on C57BL/6J). These cells are scored by subject-grouped, class-stratified k-fold cross-validation over the test-group mice, where each fold withholds a subset of the test mice, re-tunes and refits the sparse model on every remaining mouse and predictions are pooled across folds into one confusion matrix.

### 2.8 Dose response analysis using ordinal regression modeling

We then used 4 strains of mice with high withdrawal, withdrawal, and baseline observations to investigate whether the behavioral feature space encodes a graded, dose-dependent signal across escalating morphine withdrawal conditions, using a regularized ordinal regression framework.

The proportional-odds cumulative logit model was used, which assumes that an ordinal response variable Y with K ordered categories (levels of withdrawal intensity) is governed by a continuous latent variable. Following the ordinalNet [56] parameterization, the cumulative logit model is:

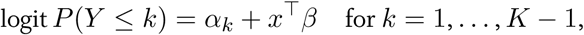

Where the feature coefficients *β* are shared across all *K−*1 cumulative logits (the proportional-odds assumption) and *α*_1_ *≤ · · · ≤ α_K−_*_1_ are ordered category thresholds. The feature matrix was *z*-scored and an *L*_1_ (lasso) penalty (*α* = 1) applied to *β*, with *λ* minimizing AIC. For visualization we defined the latent withdrawal score *η* = *−x^⊤^β*, so that higher *η* corresponds to greater withdrawal; each threshold *α_k_* is the point on this continuum where *P* (*Y ≤ k*) = 0.5.

Model classification performance was evaluated using the Ranked Probability Score (RPS) under k-fold crossvalidation. Folds were assigned at the level of the individual mouse rather than the individual observation so that no mouse contributed to both the training and held-out sets within a fold. Cross-validation used 10 folds (capped at the number of mice). On each fold a fresh ordinalNet model was fit on the training animals using the same model family and penalty as the full-data model (cumulative logit family, lasso penalty with *α* = 1, maximum 1000 outer iterations); features were z-scored using the training-fold mean and standard deviation, with the same statistics applied to the held-out fold, and the *λ* index was reselected within each fold as the minimum-AIC value on that fold’s training data (so neither the held-out observations’ scaling nor the full-data *λ* choice leaked into prediction). Each held-out observation’s predicted class was assigned as the withdrawal group with the highest predicted probability. The RPS was computed as an ordinal-aware measure of prediction quality for each held-out observation. The predicted category probabilities P(Y = k) were converted to cumulative predicted probabilities and the per-observation RPS was computed as: RPS = (1 / [K − 1]) × Σk (Fk − Ok)², where a perfect prediction yields RPS = 0. To contextualize the absolute RPS, an RPS skill score (RPSS) was calculated relative to a null reference model that predicts the marginal class distribution of the training data (estimated within each fold to avoid leakage), assigning every held-out observation that base-rate cumulative distribution. The RPSS score was then defined as: RPSS = 1 − (mean RPSmodel / mean RPSnull), where an RPSS score of 0 indicates the model performs no better than the null and 1 indicates perfect ordinal prediction.

To quantify uncertainty in the RPSS score, nonparametric bootstrap resampling was applied to the model RPS vector and the corresponding null RPS vector. In each of B = 10,000 bootstrap iterations, the same set of n observation indices was drawn with replacement and applied to both vectors simultaneously. The 2.5th and 97.5th percentiles of the 10,000 bootstrap skill scores defined the 95% confidence interval, and models were interpreted as better than the null model if this 95% confidence interval excluded 0.

### 2.9 Statistical analysis

To determine the effect of strain in delta phenotypes a linear model of each delta phenotype with strain as the predictor was fit. The normality of each model’s residuals was assessed with a Shapiro-Wilk normality test. Models with residuals that were not normally distributed were assessed with a Kruskal-Wallis rank sum test, a non-parametric alternative to an ANOVA. All reported models had non-normal residuals.

## 3 Results

### 3.1 Strain specific changes in withdrawal behavior

The eight strain comparison of withdrawal behaviors was conducted using an escalating once daily dose of morphine, culminating to 50 mg/kg, and followed by the precipitation of withdrawal using 10 mg/kg naloxone. Because baseline behavioral profiles differ across strains (Figure S1A), comparisons of strain specific withdrawal behavior rely on the delta phenotypes (withdrawal baseline behavior) to isolate withdrawal induced changes.

Across the 8 strains surveyed, we observed striking and significant strain-dependent variability in withdrawal behaviors following naloxone injection, as exemplified by 6 example phenotypes 30 min after naloxone injection (Figure 1A). These behaviors include traditional withdrawal behaviors such as jumping, locomotion, rearing, and freezing, as well as novel behaviors such as stretch attend posture and wall facing. Notably, all strains shows deviation from baseline levels in more than one behavior, and we observe an effect of strain in all phenotypes shown here (Table S4), but there is no consistent strain order in the change of these phenotypes. Consequently, inferring withdrawal severity with any one of these six behaviors alone would yield 6 different ranked lists and in cases where the direction of change is inconsistent across strain (rearing, wall facing) would be impossible to parse. Most notably, only one of the eight strains (C57BL/6J) increased jumping during withdrawal, the behavioral marker most used for cross strain comparisons. In contrast, 7 of 8 mice increased freezing time. Finally we observed distinct response profiles in 2 groups of strains, where C57BL/6J, CAST/EiJ, DBA/2J, and NOD/ShiLtJ mice exhibited decreased locomotion and supported rearing with no change in stretch attend behaviors during withdrawal and 129S1/SvImJ, A/J, AKR/J, and BALB/cJ mice show decreased in stretch attend bouts, without decreases in locomotion and rearing.

**Figure 1:**
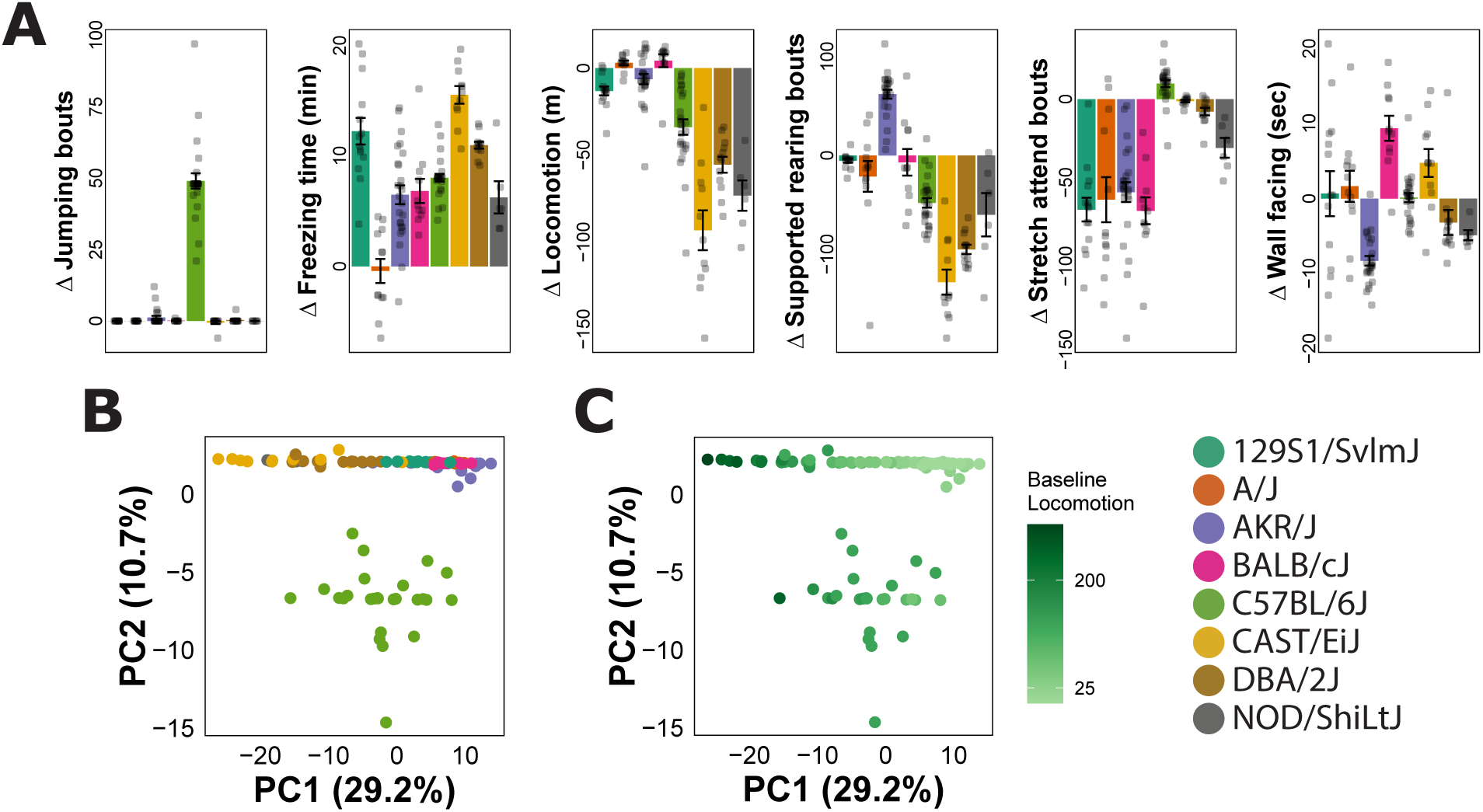
A) Changes in behavior 30 min after induction of withdrawal, where mice exhibit robust but diverse effects of withdrawal across strain (significant effect of strain, p < 0.05). B) Withdrawal induced changes in behavior identifies clustering of mice within strain and a first principal component underlain by baseline locomotion. C) Withdrawal induced changes colored by baseline locomotion at 30 min, revealing the locomotor dependent structure of PC1.

These low dimensional representations support our initial assumption that each strain has a detectable behavioral expression of withdrawal, but that expression is represented through a distinct combination of their shared behavioral repertoire. That is, there is no one globally graded effect on all behaviors across strains that might suggest a single, unified behavioral response to withdrawal across genetics. Strikingly, we only observe increases in jumping in one of eight strains, in contrast with prior results [19, 30]. This discrepancy may reflect methodological differences, including a larger testing arena or a lower naloxone dose. Regardless, the absence of elevated jumping in seven of eight strains, paired with robust alterations in other behavioral phenotypes, raises several possibilities: jumping is an inconsistent measure of opioid withdrawal across genetic backgrounds, that it is only expressed during severe opioid withdrawal, or that it is a context-specific response to withdrawal. In contrast, freezing time consistently increased across all strains except for A/J, suggesting that freezing may serve as a more stable indicator of opioid withdrawal in the open field.

Finally, the two distinct and reciprocal strain profiles of altered locomotion, supported rearing, and stretch attend suggest some underlying factor may be organizing the withdrawal response across strain. This pattern does, in fact, correspond to large differences in baseline locomotion between these two groups (Figure S1B), suggesting that general activity may influence which behavioral responses are most disrupted by withdrawal.

### 3.2 Dimensionality reduction of behavioral alterations

Given the diversity of strain-level behavioral responses to individual withdrawal behaviors, we next asked whether there was structure in the overall pattern of withdrawal-evoked behavioral changes in high dimensional behavior space. To address this, we performed sparse principal component analysis (sPCA) on the delta phenotypes across all strains using the 321 behavioral features retained from our behavioral analysis (Figure 1B). Details on sPCA model parameters and tuning can be found in table S5.

Visualization of the first two components revealed that withdrawal-induced changes clustered by strain, with most strains forming a spectrum across PC1. Further analysis revealed that, like the discrete categories observed in Figure 1A, this PC1 was organized on a spectrum based on baseline activity (Figure 1C). In contrast, PC2 distinctly segregated C57BL/6J from this PC1 spectrum.

This subsequent evidence for activity dependent withdrawal profiles along PC1 led us to hypothesize that behavioral signatures of withdrawal may interact with the activity level of a strain and we therefore defined 2 groups of strains based on their baseline locomotion (Figure S1B) to test if these intermediate groupings of strains held distinct or overlapping discriminatory information in future analyses. These groups were defined as high activity (C57BL/6J, CAST/EiJ, DBA/2J, and NOD/ShiLtJ) and low activity (129S1/SvImJ, A/J, AKR/J, BALB/cJ).

A closer inspection of PC2 loadings revealed that 15/16 of the non-zero loadings for this component were jumping related features (Figure S2), explaining the segregation of C57BL/6J mice along this component and highlighting the highly discriminative nature of jumping in detecting withdrawal in C57BL/6J mice. It also calls into question jumping’s utility as a global representation of withdrawal severity and by extension the generalization of C57BL/6J withdrawal behaviors as measures of withdrawal in other strains.

### 3.3 Indexes of withdrawal behaviors can segregate withdrawal states across genetic diversity

Having established divergent, strain-dependent patterns of withdrawal-altered behavior, we next asked whether indexes built on high dimensional behavioral data could discriminate baseline from withdrawal states using unsupervised, sPCA analysis. To address this, we conducted sPCA using all observed baseline and withdrawal trials (not delta phenotypes) in 11 groups: all mice, high activity strains, low activity strains, and within each of the 8 strains individually (Figure 2, model parameters in Table S5). When considering aggregate models, we observe that the all mice and low activity models did not clearly segregate baseline and withdrawal animals. In contrast, the high activity strain model showed modest separation of withdrawal and baseline groups. Strain-specific sPCA, however, revealed clear separation of baseline and withdrawal mice in all eight strains.

**Figure 2:**
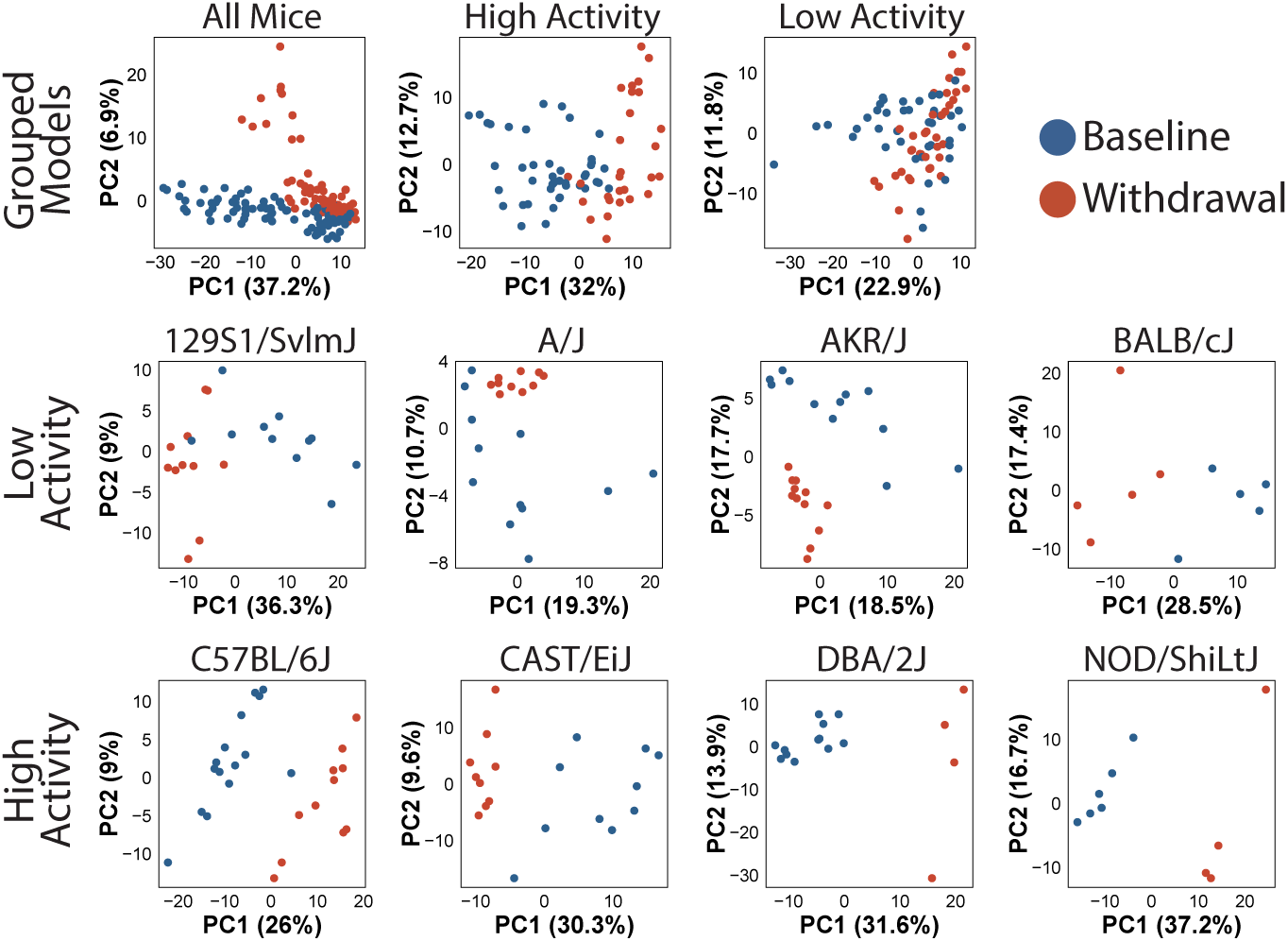
sPCA of all mice, high activity mice, low activity mice, and strain specific models. In our pooled analysis we observe separation only in the high-activity strains, suggesting they share a common response to opioid withdrawal. Further, we observe separation within each strain specific model, suggesting robust behavioral effects of withdrawal across genetics.

Lack of separation in the all mice model is likely due to the large variance in baseline behavior across strains that dominates these principal components. Interestingly, while this pattern holds true in low activity strains as well, the modest separation of withdrawal and baseline groups in the high activity model suggests a more conserved pattern of withdrawal behavior within these strains when compared to low activity strains. The clear separation observed in all 8 individual strains indicates that all 8 strains express changes in behavior following opioid withdrawal, that within-strain behavioral variation during withdrawal is sufficient for unsupervised detection of withdrawal state across genetic background, and that high-dimensional behavioral indexes are sufficient to capture that variation.

### 3.4 Supervised classification of withdrawal states across genetic backgrounds

A lack of separation in our all mice and low activity sPCA models does not necessarily indicate a lack of discriminative withdrawal features, but instead that maximal behavioral variance is explained by non-withdrawal factors. We therefore fit supervised sPLS-DA analysis to each of the 11 models to determine if and what behavioral features specifically define withdrawal. Unlike unsupervised sPCA, sPLS-DA segregated baseline and withdrawal mice in all groupings tested (Figure 3). Surprisingly, k-fold validation in all mice, high activity, and low activity models performed very robustly (accuracy = 0.93, 1, 0.9 respectively), a trend that extended into strain specific models, where accuracy ranged from 0.86-1 (Figure 4, Figure S3).

**Figure 3:**
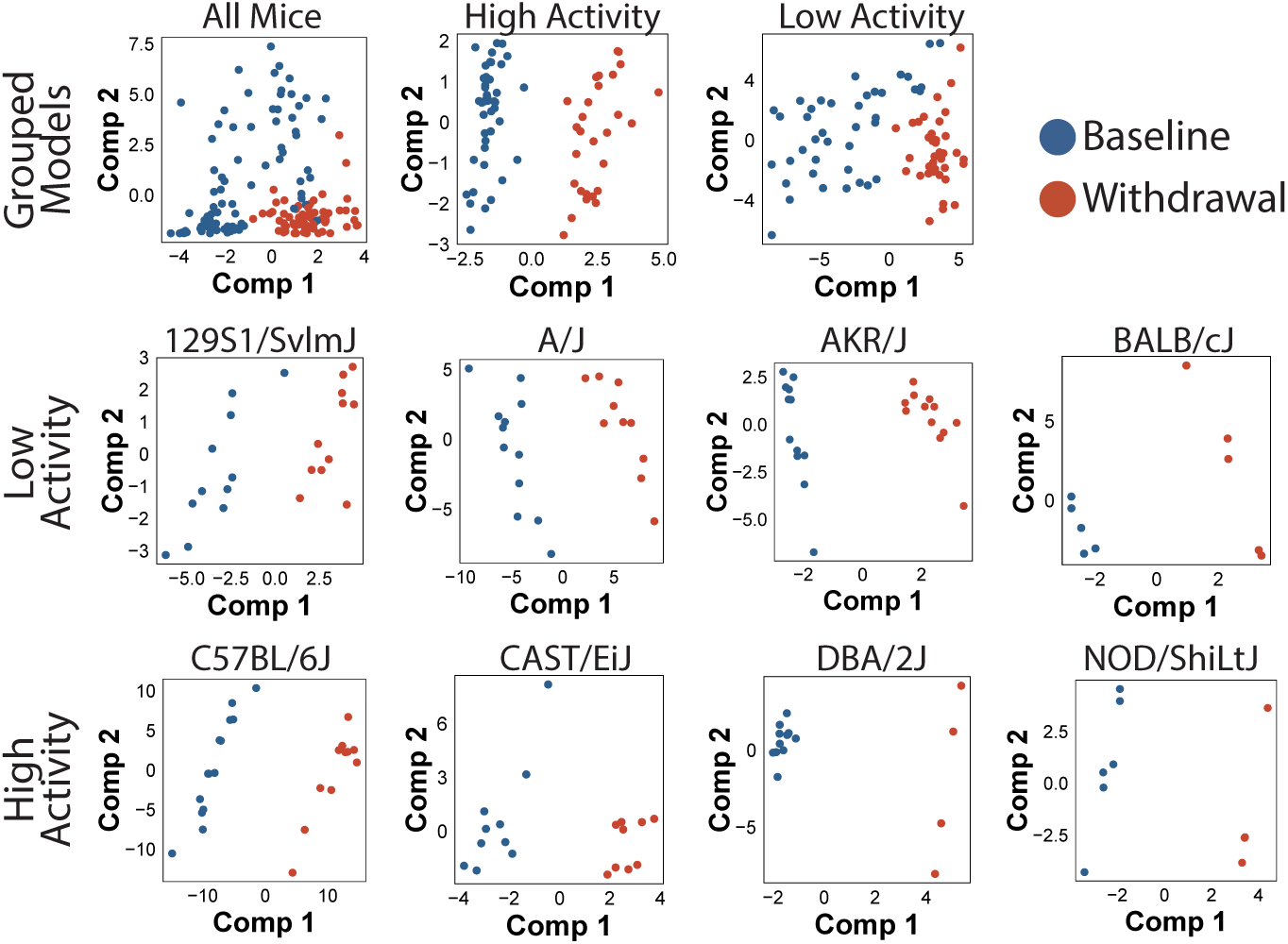
sPLS-DA of all mice, high activity mice, low activity mice, and strain specific models reveals withdrawal relevant behavioral signatures in aggregate and strain specific groups.

**Figure 4:**
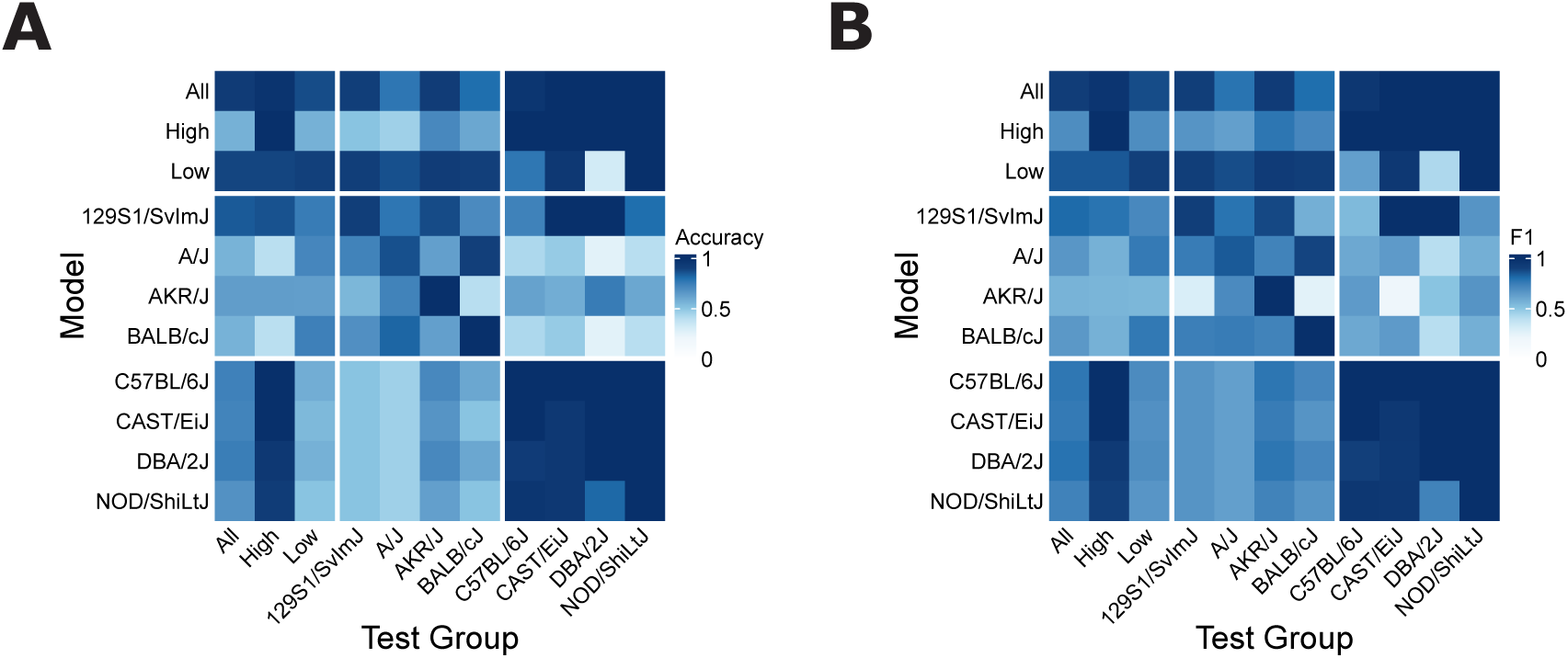
A) Within and cross model accuracy and B) F1 reveal generalizable behavior signature encoded in aggregate models, and inconsistent generalization of strain specific models across genetics.

These data suggest that aggregate models have substantial power to distinguish withdrawal and baseline behavioral profiles across genetic diversity, but that the variation defined by strain predominates this space, as observed in Figure 2. In keeping with the univariate results (Figure 1), the loadings in the all mice model include freezing on component 1, and stretch attend posture on component 2, corresponding to features altered in all mice, and low activity mice. Likewise, freezing loads onto both high and low activity sPLS-DA models, and stretch attend posture to low activity models (Supplemental materials). Unsurprisingly given these previous results, strain-specific models also robustly resolve baseline vs withdrawal states in specific backgrounds.

Robust performance within each model raised the natural next question: How well do these models generalize across strain and activity group? To answer this question we constructed models using the most generous set of features possible, and applied them to classify mice from each test set (Figure 4A, B). We observed that our all strain, high activity, and low activity models performed robustly on strains they represent (accuracy 0.86–1) with the exception of 2 comparisons (All mice model vs A/J, 0.77; All mice model vs BALB/cJ, 0.8). Interestingly, our high activity model failed to perform robustly across all low activity strains (accuracy 0.45-0.71), but low activity model accurately classified CAST/EiJ and NOD/ShLtJ mice.

When considering strain specific models we observed poor generalization across genetic diversity (strain specific model applied to all mice, accuracy = 0.56–0.84). Interestingly, strain specific models of high activity strains captured robust accuracy when applied to the high-activity test set (0.92–1), but the same was not true of low activity strains (accuracy = 0.39–0.86). These results are mirrored in the strain specific model to strain specific test set comparisons, where high activity strain models perform well on high activity strains (accuracy = 0.81-1), but not low activity strains (accuracy = 0.45-0.71) and low activity strains inconsistently generalize across both low activity (accuracy = 0.4-0.9) and high activity (0.25-1) strains.

The surprising generalization of the all mice model further reinforces the presence of some conserved, representative behavioral component of withdrawal across genetics. This is driven by high loadings of the conserved freezing features on component 1, and low-activity specific feature stretch attend posture on component 2 (Figure S4A, B), where high activity mice differ primarily on component 1, and low activity mice on a combination of component 1 and 2 (Figure S4C, D). Interestingly, the high activity strain models generalize well within high activity mice and low activity strains do not generalize within low activity, mirroring the segregation (and lack of segregation) in the high and low activity sPCAs (Figure 2) suggesting a more conserved withdrawal response in high activity mice than low activity mice.

The poor performance of strain specific models across aggregate test groups and the inconsistency of strain specific models applied to strain specific test sets underscores and confirms our original concern that strain specific models do not uniformly generalize across genetics, and is particularly salient in the low activity models, which have the worst generalization. However, this is still a concern within high activity strains, like the C57BL/6J, whose models fail to have predictive power in low activity strains.

### 3.5 Behavioral indexes can sensitively assay withdrawal intensity

Finally, for models using high dimensional behavioral data to be maximally valuable they must discriminate withdrawal intensity, because this resolution will allow hypothesis testing concerning the impact of genetic variation or pharmacology on withdrawal intensity. We therefore used ordinal regression models to classify baseline, withdrawal, and high withdrawal mice in 4 distinct genetic backgrounds (129S1/SvImJ, BALB/cJ, C57BL/6J, and DBA/2J), and within an aggregate model representing all 4 strains (Figure 5). We observed the 129S1/SvImJ, C57BL/6J, and DBA/2J models predict dose significantly better than null models (respectively, RPSS = 0.34, 0.76, 0.55; overall accuracy = 0.64, 0.83, 0.82), but not the BALB/cJ model (Table S7). Excitingly, when predicting withdrawal group in the four strain model we observed robust classification performance (RPSS = 0.62).

**Figure 5:**
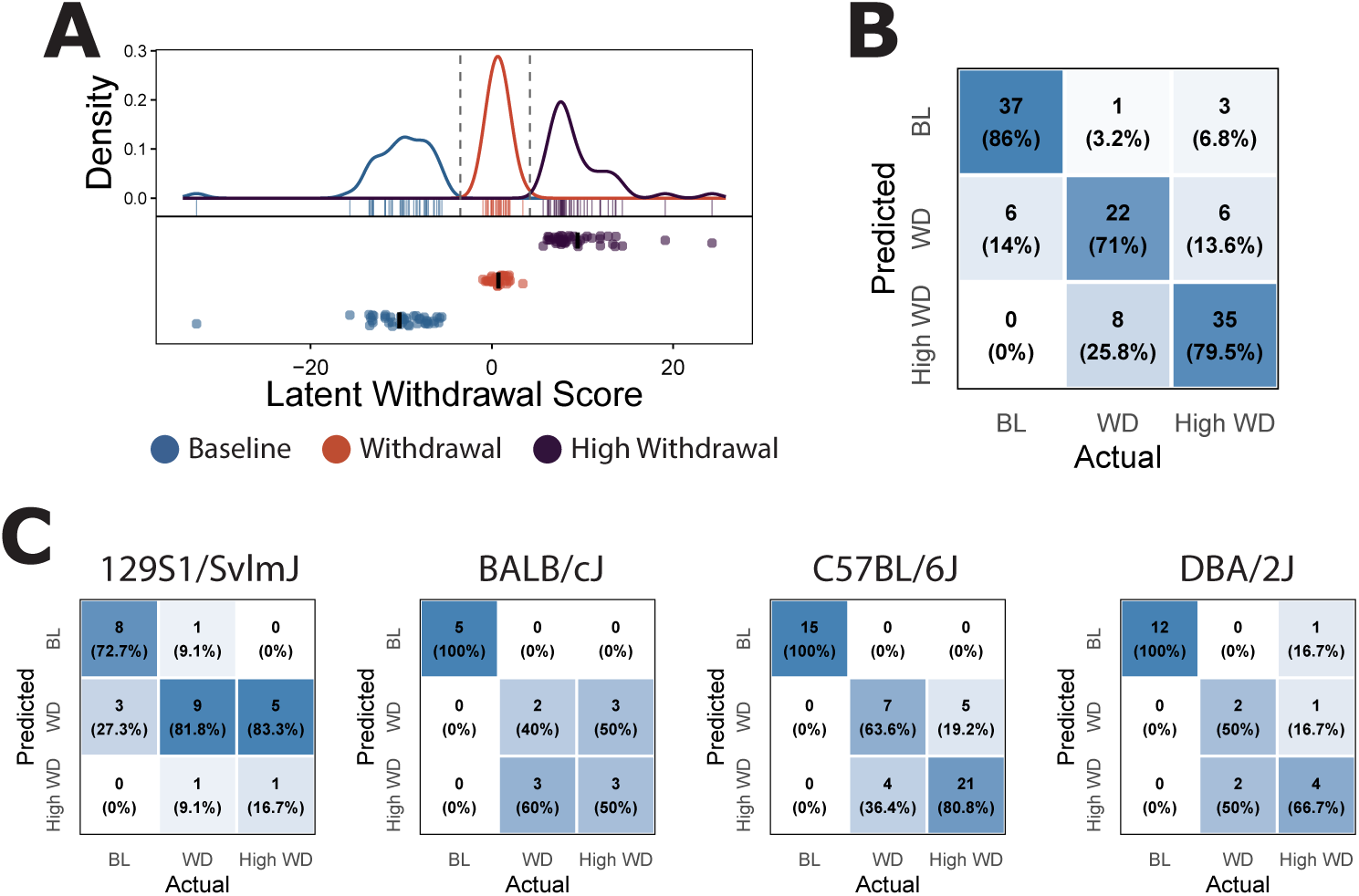
A) Ordinal regression modeling successfully classifies withdrawal group significantly better than the null model, revealing behavior signature associated with withdrawal severity. B) Confusion matrix of the four strain model of withdrawal severity. C) Strain specific modeling of withdrawal severity successfully discriminates opioid withdrawal severity in 129S1/SvImJ, C57BL/6J, DBA/2J, but not BALB/cJ mice.

These results support the use of high dimensional behavior analysis in predicting withdrawal severity within an individual strain, and hold promise for investigations into the genetic, neural, and pharmacological mediators of withdrawal. While discrimination of withdrawal groups within the all model is exciting, it is important to remember that this model is fit to withdrawal group, not withdrawal intensity per se. While it is reasonable to assume that escalating morphine exposure grades withdrawal intensity within a strain, there is no way to confirm the relative withdrawal intensity at a dose of morphine and naloxone across strains. However, this does extend our prior observation that core withdrawal features may, in the future, be leveraged across genetics to study withdrawal severity. In congruence with our binary classification, the most informative features in this all strains model are related to freezing time, and number of freezing bouts (Figure S5).

## 4 Discussion

We demonstrate that the behavioral signatures of opioid withdrawal can be captured using computer vision and machine learning based analysis, and that indexes built on high-dimensional behavioral data reliably encode withdrawal within individual strains and in genetically aggregated groups. Further, they encode graded withdrawal within three of the four strains tested. The robust, but strain-specific, discrimination of withdrawal is well suited to screening interventions within inbred strains, accelerating the preclinical study of withdrawal. Finally, our results argue that single behaviors are unreliable proxies for the latent state of withdrawal, and that comparing any one behavior across strains risks comparing the biology that maps the latent state of withdrawal onto observable behavior rather than the severity of the clinically relevant latent state itself.

It is only through automated computer vision that we could sample a behavioral space large enough to predict withdrawal across diverse genetics with meaningful throughput. JABS provided access to a behavioral feature space too large to score by hand (321–652 features/model), preserved the temporal and spatial structure of behavior, and captured measures like gait dynamics and spinal posture that human observers cannot feasibly score at scale. This approach not only enriches the depth of behavioral phenotypes, but pushes preclinical screening towards the multidimensional outcomes and assessments seen in clinical practice (e.g., the Clinical Opiate Withdrawal Scale [54]). This approach, therefore, seems well suited for other complex, multifactorial, latent states relevant to neuropsychiatric illnesses like pain, depression, or developmental disorders.

### 4.1 High dimensional behavioral indexes encode robust, but not universally transferable, signatures of withdrawal within genetic backgrounds

The most fundamental question answered in this work was to what extent high-dimensional behavioral indexes can resolve withdrawal across genetic diversity. Our approach discerned opioid withdrawal in every strain surveyed: strain-specific sPCA separated baseline from withdrawal mice in all eight strains, and strain-specific sPLSDA classified withdrawal with cross-validated accuracy of 0.86–1.00. These results support the sensitivity of our approach and the assertion that all genetic backgrounds experienced some form of latent withdrawal state. However, while withdrawal was detectable in all tested strains, the behavioral encoding of withdrawal was not interchangeable across genotypes, and strain-specific models generalized poorly and inconsistently outside their own activity group (Figure 4). The C57BL/6J model, for example, perfectly classified the other three high-activity strains (CAST/EiJ, DBA/2J, and NOD/ShiLtJ), yet performed poorly on each low-activity strain (accuracy = 0.45– 0.71) and on the pooled test set of all other strains (accuracy = 0.73). It is therefore dangerous to generalize any withdrawal index constructed on a single genetic background to diverse populations, as it risks misrepresenting withdrawal severity in diverse genetics by ignoring the influence of genetics on the behavioral expression of withdrawal.

However, this represents an unrealized opportunity. Because the mapping from latent withdrawal state to observable behavior is comparatively stable within a fixed background (demonstrated by our within-model crossvalidation), these strain-specific models represent sensitive assays for perturbations within each background, and a panel of such models spans genetic diversity. This framework is well suited to genetic, neural, and pharmacological screening, where single-gene knockouts can be compared against isogenic wild-type controls; circuit-level manipulations can be read out against the same stable signature; and candidate therapeutics can be evaluated across multiple backgrounds, each scored by its own validated model, to determine whether efficacy is genotype-dependent. This opportunity is made more salient by the extension of these strain-specific signatures to graded predictions. Because the severity of opioid withdrawal scales with the magnitude of prior opioid exposure in mice and rats [10, 36], escalating morphine exposure provides an experimentally controlled, ordinal manipulation of dependence depth within a background, against which candidate measures can be evaluated. Three of four strain-specific ordinal models (129S1/SvImJ, C57BL/6J, and DBA/2J) predicted withdrawal group (baseline, withdrawal, high withdrawal) better than a base-rate null (RPSS 0.34–0.76), whereas the BALB/cJ model did not. These results establish high-dimensional behavior indexes not just as a detector of withdrawal but as a sensitive, severity-graded read-out within a genetic background, suitable for hypothesis-driven study of genetic, neural, and pharmacological modifiers of withdrawal.

### 4.2 High dimensional behavioral indexes detect a partially conserved signature of withdrawal across genetic backgrounds

The lack of transferability between strain-specific models naturally raises our next question: is there a conserved behavioral signature of withdrawal across genetic diversity? Unsupervised sPCA did not clearly separate baseline from withdrawal mice in the all-mouse or low-activity groups, likely because variance in baseline behavior across strains dominated the leading components. Supervised sPLS-DA, however, recovered a withdrawal axis in every aggregated group, with cross-validated accuracy of 0.93 (all mice), 1.00 (high activity), and 0.90 (low activity), supporting the assertion that withdrawal can, in some way, be assayed in a genetically agnostic manner. The all-mouse model also classified the mice of each individual strain with accuracy of 0.77–1.00. Its first component contrasted freezing, which increased during withdrawal in seven of eight strains in univariate analysis, with wall facing, and its second component was defined by stretch attend posture, which decreased during withdrawal primarily in low-activity strains. Most impactfully, this observation extended into withdrawal severity prediction, where an aggregate ordinal model pooling the four dose-response strains performed comparably to the strain-specific models (RPSS 0.62), with freezing time and freezing bouts again among its most heavily weighted features.

While exciting, this observation comes with caveats, primarily that the ability to predict an ordinally defined withdrawal condition across genetics does not necessarily correspond with the ability to predict the latent affective state accompanying withdrawal. That is, mice of different strains receiving the same opioid and naloxone exposure may experience different withdrawal intensities that share some mutual behavioral signature. Within a single inbred strain, where the expression of withdrawal behavior can be assumed stable, a feature set that tracks exposure is sensitive to graded withdrawal in that background. Across backgrounds, such sensitivity is a necessary step toward, but not equivalent to, a severity scale calibrated across genetic diversity. Therefore, future work tying shared behavioral facets of withdrawal responses to physiological markers of withdrawal is needed to make genetically agnostic models predictive of latent withdrawal states instead of doses.

### 4.3 Baseline activity partially organizes the behavioral expression of withdrawal

Baseline activity emerged as one organizing axis of withdrawal behavior across strains. The four strains with high baseline locomotion (C57BL/6J, CAST/EiJ, DBA/2J, and NOD/ShiLtJ) decreased locomotion and supported rearing during withdrawal with no change in stretch attend posture, whereas the four low-activity strains (129S1/SvImJ, A/J, AKR/J, and BALB/cJ) decreased stretch attend bouts without a decrease in locomotion and rearing. The first principal component of withdrawal-induced behavioral change likewise ordered strains along a spectrum of baseline locomotion. The withdrawal response also appeared more conserved within the high-activity group, and high-activity strain models transferred well to the other high-activity strains (accuracy = 0.81–1.00), whereas low-activity strain models transferred inconsistently to the other low-activity strains (accuracy = 0.40–0.90).

Failures of the high-activity strain models on low-activity strains were almost entirely false positives. Each of these models retained perfect sensitivity on every low-activity strain while specificity fell to 0.00–0.42 (Figure S3), meaning that they classified most low-activity baseline mice as in withdrawal. This pattern is expected if reduced locomotion and rearing, prominent features of withdrawal in high-activity strains, resemble the baseline state of low-activity strains, and it illustrates how a genotype’s baseline propensity to perform a behavior can be mistaken for withdrawal. The groups were defined from this cohort, however, so their generality remains to be tested across new cohorts and a wider selection of genetic diversity.

### 4.4 Single behaviors are unreliable proxies for the latent state of withdrawal

Most obviously, our results contend that no single behavior can index the severity of a complex, latent state of withdrawal across genetics. Behaviors that we observe during withdrawal are filtered through each genotype’s behavioral repertoire and therefore any single behavior risks conflating the severity of withdrawal and the genotype-dependent propensity to perform that behavior. The six representative behaviors we examined in univariate analysis illustrate this: each showed a significant effect of strain, yet each would produce a different ranking of strain withdrawal severity, and two of them (supported rearing and wall facing) changed in opposing directions in different strains.

Jumping serves as a particularly salient, historically relevant example. In this work only one of eight strains (C57BL/6J) increased jumping during withdrawal. The remaining seven strains showed no increase in jumping, yet the withdrawal state of each was classified with high accuracy by its own strain-specific model. Estimating withdrawal from jumping alone would therefore misrepresent the latent state in most of the genotypes we examined and, by extension, would distort severity comparisons among them. This result bears directly on prior strain surveys of withdrawal that relied on jumping [19, 30], in which jumping was not genetically correlated with other withdrawal signs such as hypothermia or fecal boli production [30], and on whether such surveys measured genetic predisposition to withdrawal severity or genetic propensity to jump during withdrawal. Compared with those studies, we used a larger testing arena and a lower naloxone dose, either of which could account for the discrepancy in jumping between our observations and theirs.

Freezing was the most consistent single behavior we measured, increasing in seven of eight strains and carrying substantial weight in the pooled classification and severity models. Even freezing, however, did not increase in A/J mice, whose withdrawal was nonetheless detected by their strain-specific model (accuracy = 0.86). Our data therefore show that single behaviors cannot reliably assess withdrawal across genetic diversity.

### 4.5 Limitations

Several limitations qualify these conclusions. Sample sizes were modest for several strains (e.g., NOD/ShiLtJ, n = 6; BALB/cJ, n = 11), limiting the stability of strain-specific embedding spaces. This concern extends to the BALB/cJ severity model that did not outperform the null (n = 11 mice, 16 observations). Sex balance varied across strain and treatment groups, and sex was not modeled as a factor, so these analyses cannot resolve sex differences in the behavioral expression of withdrawal. Mice tested in both baseline and withdrawal conditions encountered the arena twice, and although baseline testing preceded the first morphine injection by 3–4 days, prior exposure cannot be fully excluded as an influence on withdrawal behavior. Because not all mice were tested in both conditions, delta phenotypes for mice lacking one observation were completed with the median of their strainby-treatment group, and in the standard withdrawal arm no AKR/J, BALB/cJ, or C57BL/6J mouse contributed a within-subject pair; delta phenotypes for these strains therefore reflect withdrawal relative to a strain median baseline. Finally, the high- and low-activity groups were defined after inspecting withdrawal-induced change in this same cohort, and each contains four strains, so their value as a general organizing principle requires testing in strains not used to define them.

### 4.6 Future Directions

Several natural extensions of this work would strengthen and generalize these findings. Connecting behavioral classification to physiological measures of withdrawal would provide critical validation and mechanistic insight for both strain-specific and genetically agnostic models. Correlating model-predicted withdrawal severity with circuit-level activation markers (e.g., cFos mapping), stress axis engagement (e.g., corticosterone), and molecular signatures of withdrawal (e.g., cAMP rebound) would validate the behavioral models and identify behavioral markers that represent a conserved expression of withdrawal across genetics.

Expansion to larger and more genetically diverse mouse panels, including the BXD recombinant inbred strains, the Collaborative Cross, and the Diversity Outbred population, would test whether the activity-level grouping generalizes beyond the eight inbred strains used to define it. The Diversity Outbred population is particularly relevant because each animal is genetically unique, providing a direct test of whether activity-based classification extends to continuous genetic diversity. These larger panels could also enable genetic mapping using the expanded behavioral phenotypes developed here, potentially identifying loci associated with specific behavioral dimensions of withdrawal rather than a single composite score.

Finally, extending automated phenotyping to 24-hour monitoring of spontaneous withdrawal would improve ecological validity and avoid both the pharmacological induction of withdrawal by naloxone and any genetic effects specific to naloxone pharmacology. We used precipitated withdrawal here because the effect of genetics on the kinetics of spontaneous withdrawal is unknown; continuous monitoring would measure those kinetics directly in each strain. Continuous monitoring would also give access to circadian patterns of withdrawal behavior, as shown in rats [31], withdrawal during the dark (active) phase, and social context [17], all of which are translationally relevant considerations.

### 4.7 Conclusions

Opioid withdrawal behavior is profoundly shaped by genetic background, and investigations into genetic, pharmacological, or neural mediators of opioid withdrawal would benefit greatly from high-dimensional genotype-informed behavioral models. Combining automated computer-vision phenotyping with genotype-aware multivariate modeling provides a scalable path toward genetically informed withdrawal assessment in diverse mouse populations, with the ultimate goal of identifying the genetic and biological networks that modulate opioid withdrawal and may represent new targets for treating opioid use disorder.

## Supporting information

Feature definitions and model loadings

## 5 Acknowledgments

We thank Kumar Lab members for their help and comments. This work was funded by The Jackson Laboratory Directors Innovation Fund, National Institutes of Health, National Institute on Drug Abuse DA051235, DA050837 (V.K.), National Institute on Aging AG078530 (V.K.), and National Institute on Mental Health MH138309 (V.K.).

## 6 Supplement

### 6.1 Supplemental Figures

**Figure S1:**
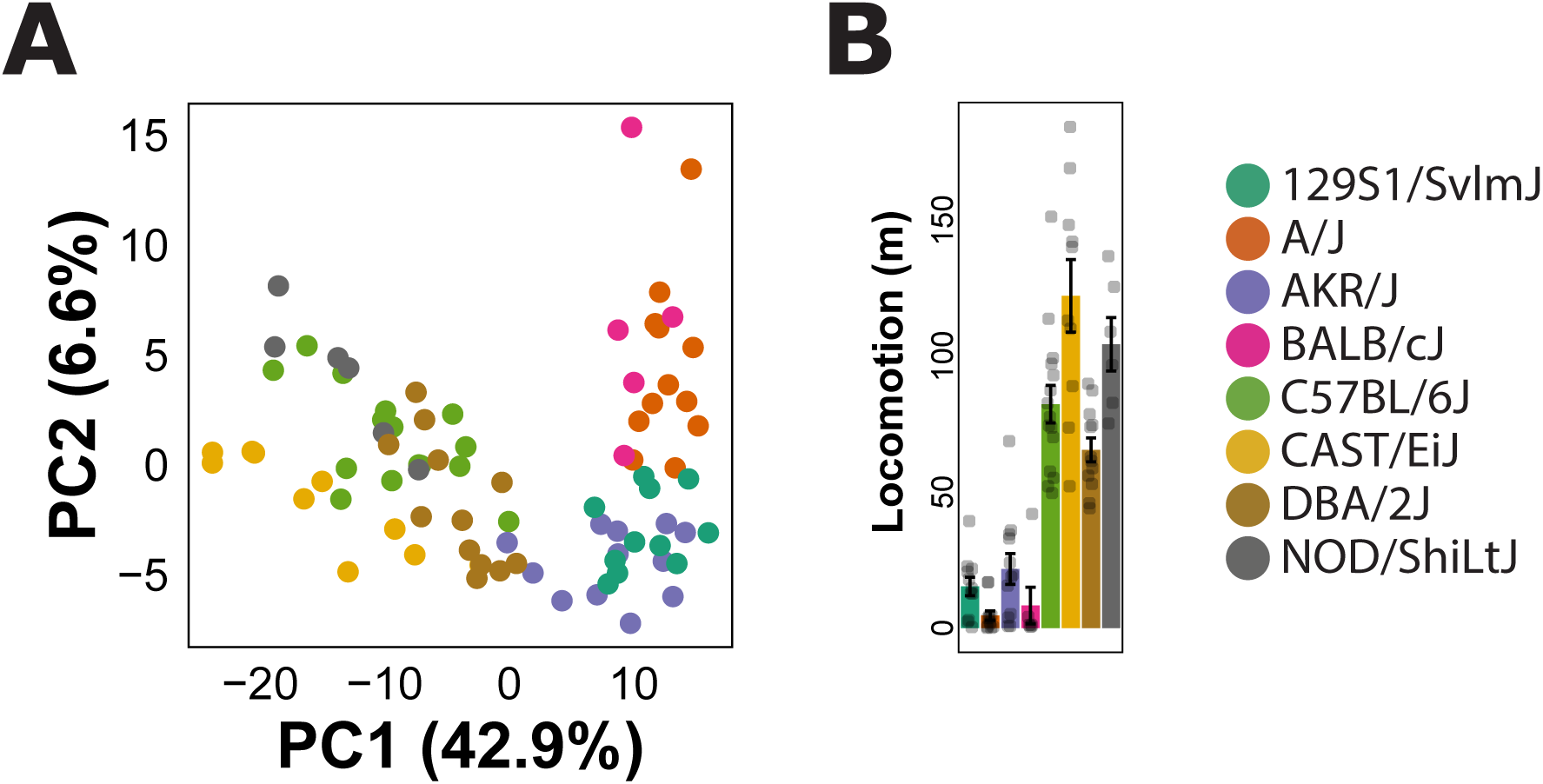
A) sPCA analysis of all baseline mice revealing clustering of behavioral profiles based on strain. B) Baseline locomotion at 30 mins, revealing 4 extremely low activity strains.

**Figure S2:**
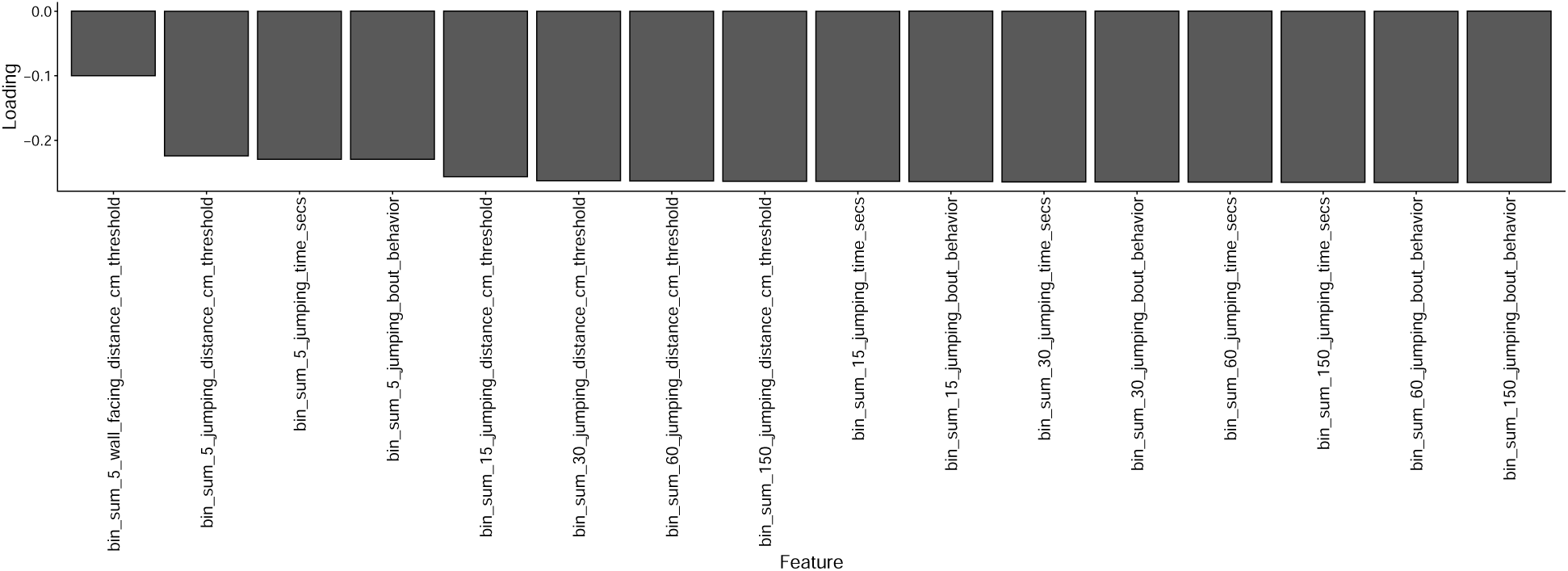
Non-zero feature loadings for PC2 of delta phenotype sPCA.

**Figure S3:**
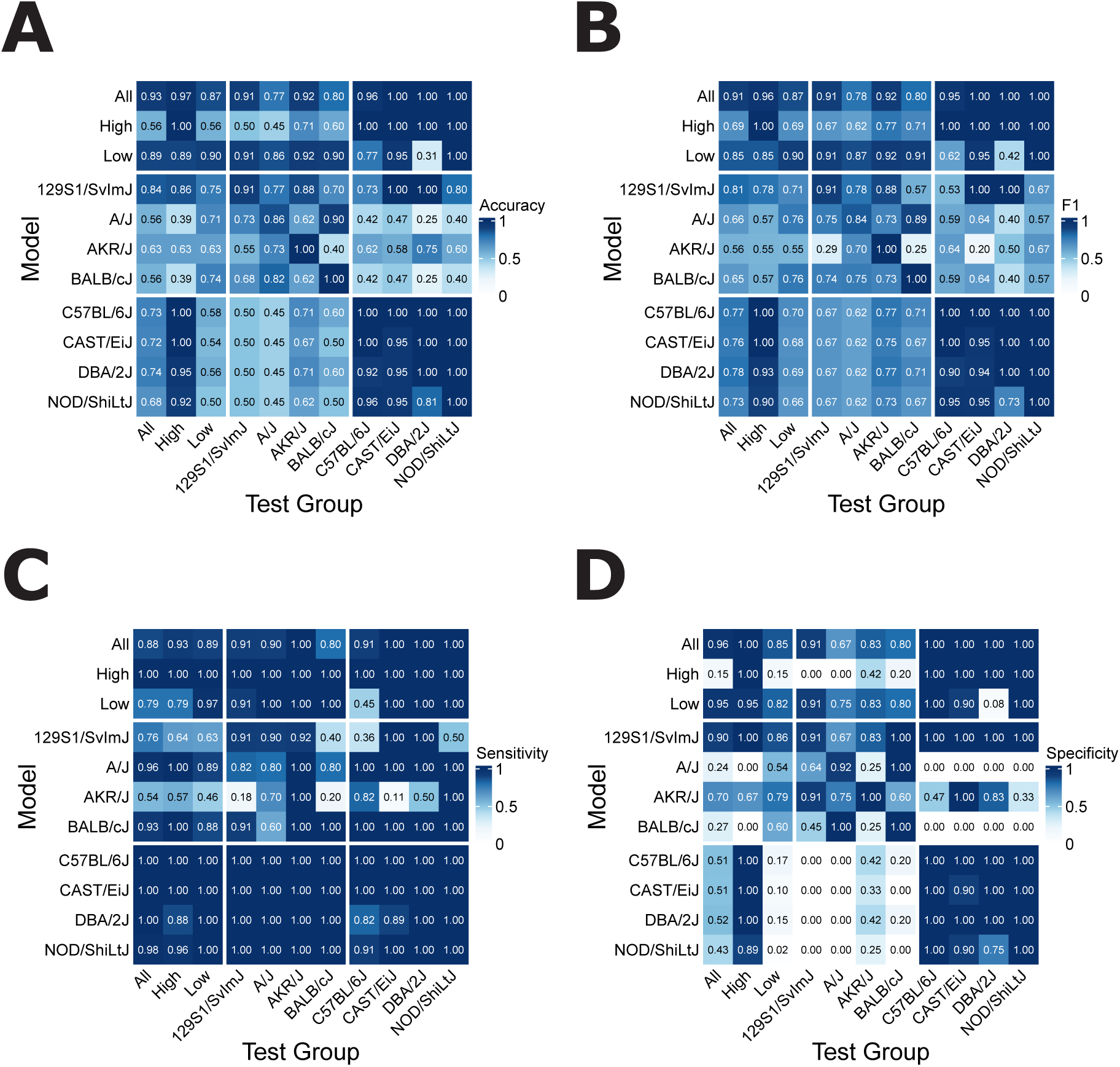
A) Accuracy B) F1 Beta C) Sensitivity and D) Specificity of within and cross model performance

**Figure S4:**
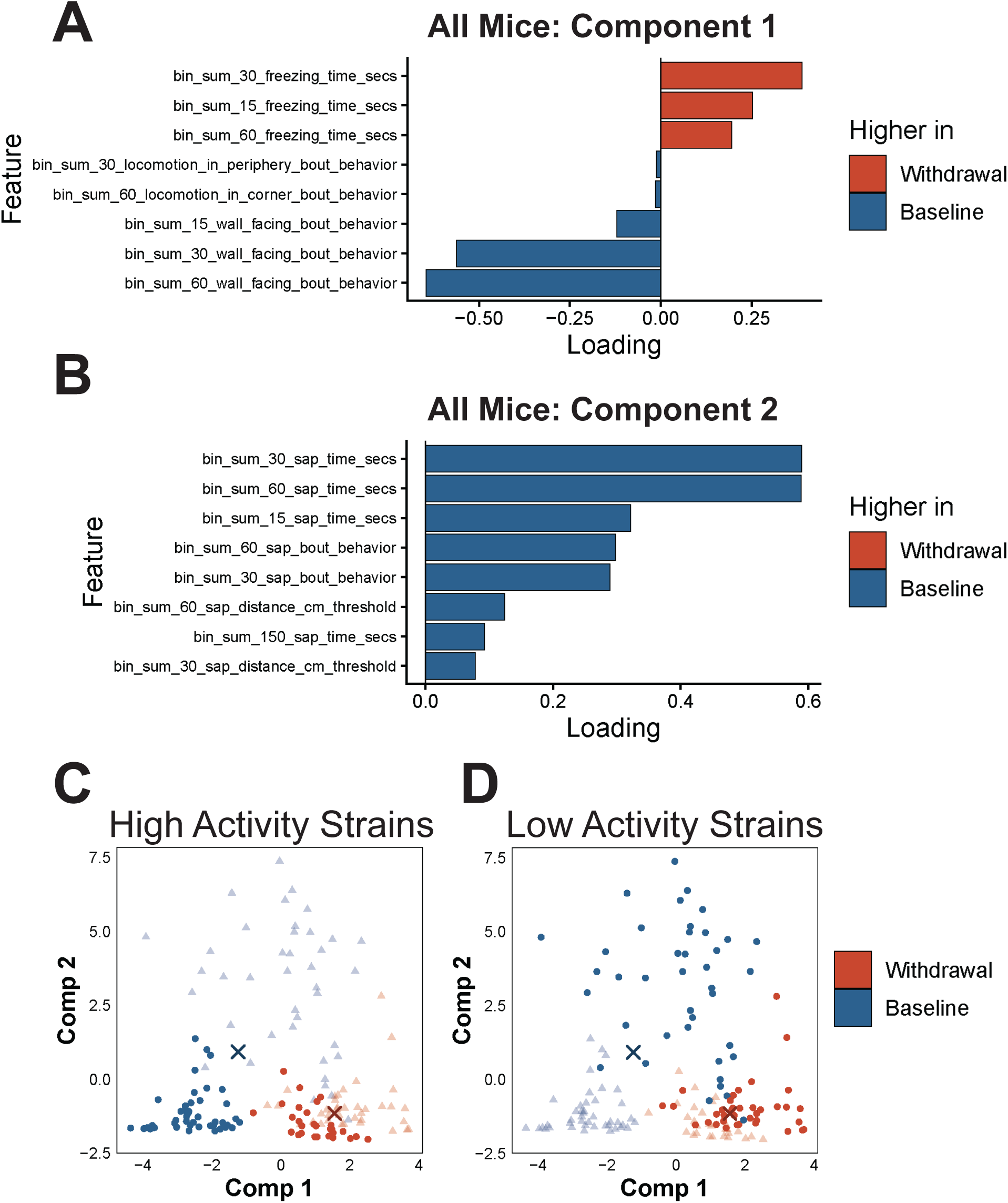
Feature loadings for all mice sPLS-DA model A) component 1 B) component 2. sPLS-DA all mice model with C) high-activity mice and D) low-activity mice highlighted in the embedding space. Dark circles represent mice within the selected group, transparent triangles represent mice of the opposite group. X’s represent centroids for each class.

**Figure S5:**
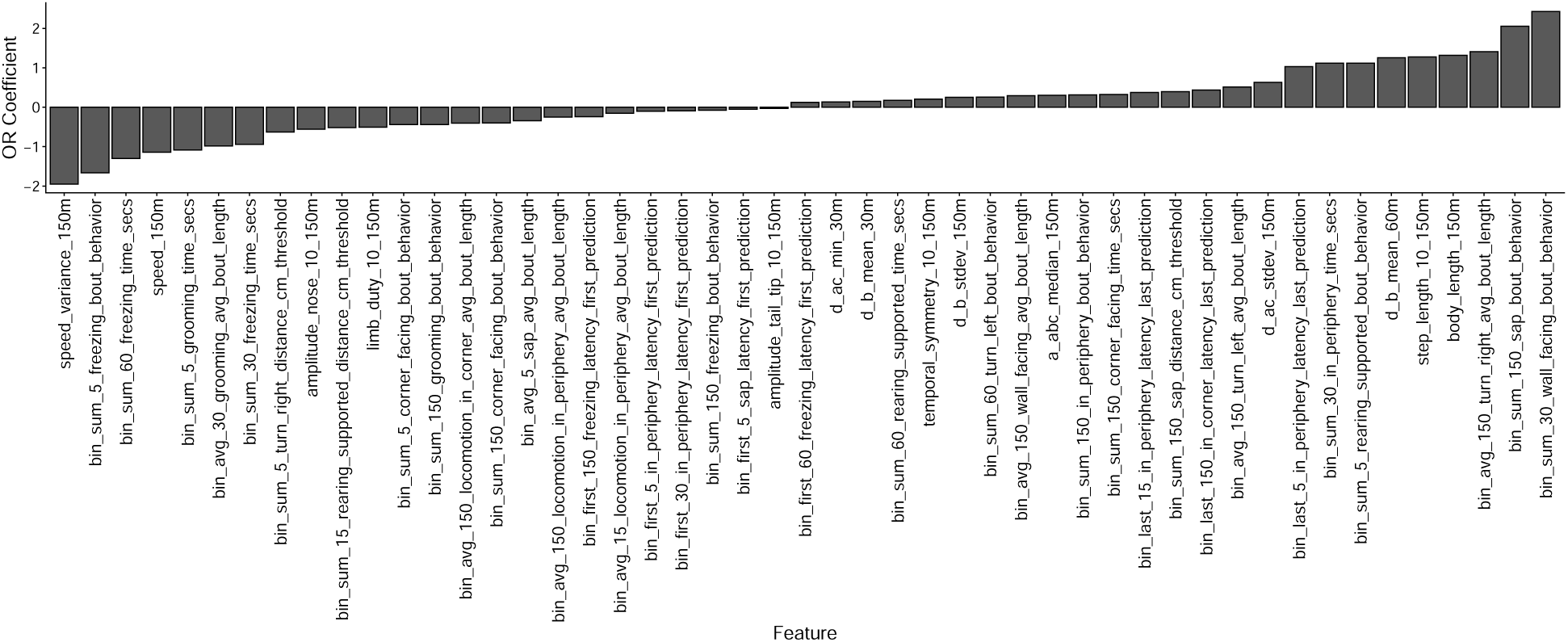
Coefficients for four strain ordinal regression model showing high loadings for conserved freezing features

### 6.2 Supplemental Tables

**Table S1:** Sample size.

| Strain | JAX ID | Baseline Only | Withdrawal Only | Both | High Withdrawal Only | High Both | Total |
| --- | --- | --- | --- | --- | --- | --- | --- |
| 129S1/SvImJ | 002448 | 0 | 3 | 8 | 3 | 3 | 17 |
| A/J | 000646 | 2 | 0 | 10 | 0 | 0 | 12 |
| AKR/J | 000648 | 12 | 12 | 0 | 0 | 0 | 24 |
| BALB/cJ | 000651 | 0 | 5 | 0 | 1 | 5 | 11 |
| C57BL/6J | 000664 | 12 | 11 | 0 | 23 | 3 | 49 |
| CAST/EiJ | 000928 | 2 | 1 | 8 | 0 | 0 | 11 |
| DBA/2J | 000671 | 2 | 0 | 4 | 0 | 6 | 12 |
| NOD/ShiLtJ | 001976 | 2 | 0 | 4 | 0 | 0 | 6 |
| High Activity | NA | 18 | 12 | 16 | 23 | 9 | 78 |
| Low Activity | NA | 14 | 20 | 18 | 4 | 8 | 64 |
| All mice | NA | 32 | 32 | 34 | 27 | 17 | 142 |

**Table S2:** Sample size by sex.

| Strain | Sex | JAX ID | Baseline Only | Withdrawal Only | Both | High Withdrawal Only | High Both | Total |
| --- | --- | --- | --- | --- | --- | --- | --- | --- |
| 129S1/SvImJ | F | 002448 | 0 | 3 | 2 | 3 | 0 | 8 |
| 129S1/SvImJ | M | 002448 | 0 | 0 | 6 | 0 | 3 | 9 |
| A/J | F | 000646 | 1 | 0 | 5 | 0 | 0 | 6 |
| A/J | M | 000646 | 1 | 0 | 5 | 0 | 0 | 6 |
| AKR/J | F | 000648 | 7 | 6 | 0 | 0 | 0 | 13 |
| AKR/J | M | 000648 | 5 | 6 | 0 | 0 | 0 | 11 |
| BALB/cJ | F | 000651 | 0 | 2 | 0 | 0 | 3 | 5 |
| BALB/cJ | M | 000651 | 0 | 3 | 0 | 1 | 2 | 6 |
| C57BL/6J | F | 000664 | 6 | 5 | 0 | 11 | 0 | 22 |
| C57BL/6J | M | 000664 | 6 | 6 | 0 | 12 | 3 | 27 |
| CAST/EiJ | F | 000928 | 0 | 0 | 6 | 0 | 0 | 6 |
| CAST/EiJ | M | 000928 | 2 | 1 | 2 | 0 | 0 | 5 |
| DBA/2J | F | 000671 | 0 | 0 | 0 | 0 | 3 | 3 |
| DBA/2J | M | 000671 | 2 | 0 | 4 | 0 | 3 | 9 |
| NOD/ShiLtJ | M | 001976 | 2 | 0 | 4 | 0 | 0 | 6 |
| High Activity | F | NA | 6 | 5 | 6 | 11 | 3 | 31 |
| High Activity | M | NA | 12 | 7 | 10 | 12 | 6 | 47 |
| Low Activity | F | NA | 8 | 11 | 7 | 3 | 3 | 32 |
| Low Activity | M | NA | 6 | 9 | 11 | 1 | 5 | 32 |
| All mice | F | NA | 14 | 16 | 13 | 14 | 6 | 63 |
| All mice | M | NA | 18 | 16 | 21 | 13 | 11 | 79 |

**Table S3:** Validation performance of freezing and jumping classifiers.

| Classifier | Videos | Behavior frames | Not-behav. frames | k-fold F1 | k-fold Accuracy |
| --- | --- | --- | --- | --- | --- |
| Jumping | 18 | 3230 | 8890 | 0.96 | 0.98 |
| Freezing | 22 | 18380 | 15047 | 0.95 | 0.97 |

**Table S4:** Significance testing for effect of strain in individual phenotypes.

| Phenotype | Shapiro-Wilk p-val | Test | $\chi^2$ | df | Effect of strain; Kruskal-Wallis p |
| --- | --- | --- | --- | --- | --- |
| bin_sum_30_jumping_bout_behavior | 7.2e-19 | Kruskal-Wallis | 95.1 | 7 | 1.1e-17 |
| bin_sum_30_freezing_time_secs | 1.3e-2 | Kruskal-Wallis | 68.1 | 7 | 3.6e-12 |
| bin_sum_30_locomotion_distance_cm_threshold | 1.5e-5 | Kruskal-Wallis | 86 | 7 | 8.1e-16 |
| bin_sum_30_sap_bout_behavior | 5.0e-7 | Kruskal-Wallis | 82.2 | 7 | 4.8e-15 |
| bin_sum_30_wall_facing_time_secs | 7.0e-6 | Kruskal-Wallis | 60.5 | 7 | 1.2e-10 |
| bin_sum_30_rearing_supported_bout_behavior | 3.0e-5 | Kruskal-Wallis | 90.9 | 7 | 8.3e-17 |

**Table S5:** Tuning and final model parameters for the sparse PCA models.

| Model | Folds | Fold sizes | Repeats | Null correlation | Components searched | Components fitted | Proportion of retained features | Nonzero loadings per component | Cumulative variance explained |
| --- | --- | --- | --- | --- | --- | --- | --- | --- | --- |
| all_mice_baseline | 5 | 17/17/17/16/16 | 5 | 0.205 | 12 | 12 | 0.8013/0.5994/<br>0.4006/0.5994/<br>0.8013/0.8013/<br>0.1987/0.1987/<br>0.8013/0.4006/<br>0.4006/0.1987 | 254/190/127/<br>190/254/254/<br>63/63/254/<br>127/127/63 | 0.7820 |
| all_mice | 5 | 30/30/30/30/29 | 5 | 0.150 | 14 | 14 | 0.6012/0.8006/<br>0.0498/0.1994/<br>0.0249/0.6012/<br>0.0997/0.0498/<br>0.0997/0.3988/<br>0.1994/0.0249/<br>0.0249/0.8006 | 193/257/16/<br>64/8/193/32/<br>16/32/128/64/<br>8/8/257 | 0.7619 |
| C57BL6j | 5 | 6/5/5/5/5 | 5 | 0.415 | 12 | 4 | 0.5997/0.4003/<br>0.8006/0.8006 | 391/261/522/<br>522 | 0.4897 |
| AKRJ | 5 | 5/5/5/5/4 | 5 | 0.440 | 11 | 4 | 0.099/0.099/0.6/<br>0.8 | 52/52/315/<br>420 | 0.5057 |
| 129S1SvImJ | 5 | 5/5/4/4/4 | 5 | 0.470 | 9 | 3 | 0.8/0.6/0.0514 | 280/210/18 | 0.5070 |
| CASTeij | 5 | 4/4/4/4/3 | 5 | 0.527 | 9 | 3 | 0.1997/0.8003/<br>0.3993 | 122/489/244 | 0.4793 |
| DBA2j | 5 | 4/3/3/3/3 | 5 | 0.609 | 7 | 5 | 0.4003/0.8006/<br>0.8006/0.0997/<br>0.0249 | 257/514/514/<br>64/16 | 0.6786 |
| NODShiLtj | 3 | 4/3/3 | 5 | 0.591 | 5 | 2 | 0.6006/0.3994 | 382/254 | 0.5383 |
| AJ | 5 | 5/5/4/4/4 | 5 | 0.470 | 10 | 3 | 0.2/0.0493/<br>0.0493 | 73/18/18 | 0.3830 |
| BALBcJ | 3 | 4/3/3 | 5 | 0.591 | 5 | 2 | 0.8009/0.8009 | 354/354 | 0.4595 |
| deltas | 5 | 23/23/23/23/23 | 5 | 0.172 | 15 | 15 | 0.6012/0.0498/<br>0.0249/0.3988/<br>0.8006/0.3988/<br>0.0997/0.3988/<br>0.3988/0.3988/<br>0.3988/0.3988/<br>0.8006/0.0498/<br>0.3988 | 193/16/8/128/<br>257/128/32/<br>128/128/128/<br>128/128/257/<br>16/128 | 0.7684 |
| high_activity | 5 | 15/14/14/14/14 | 5 | 0.224 | 11 | 10 | 0.3988/0.6012/<br>0.1994/0.0249/<br>0.1994/0.1994/<br>0.0249/0.0249/<br>0.0249/0.3988 | 128/193/64/8/<br>64/64/8/8/8/<br>128 | 0.7397 |
| low_activity | 5 | 16/16/16/15/15 | 5 | 0.213 | 16 | 13 | 0.6012/0.8006/<br>0.6012/0.8006/<br>0.0249/0.6012/<br>0.8006/0.0498/<br>0.3988/0.0249/<br>0.0498/0.0997/<br>0.6012 | 193/257/193/<br>257/8/193/<br>257/16/128/8/<br>16/32/193 | 0.7178 |

**Table S6:** Tuning and final model parameters for the sparse PLS-DA models.

| Model | Observations | Features | Components fitted | Components selected by CV | Nonzero loadings per component |
| --- | --- | --- | --- | --- | --- |
| all_mice | 149 | 321 | 3 | 3 | 8\|8\|32 |
| C57BL6J | 26 | 652 | 2 | 1 | 261\|130 |
| AKRJ | 24 | 525 | 2 | 1 | 13\|13 |
| 129S1SvImJ | 22 | 350 | 2 | 2 | 18\|9 |
| CASTeij | 19 | 611 | 2 | 1 | 15\|15 |
| DBA2J | 16 | 642 | 2 | 1 | 16\|16 |
| NODShiLtJ | 10 | 636 | 2 | 1 | 16\|16 |
| AJ | 22 | 365 | 2 | 1 | 146\|146 |
| BALBcJ | 10 | 442 | 2 | 1 | 11\|44 |
| high_activity | 71 | 321 | 2 | 1 | 8\|8 |
| low_activity | 78 | 321 | 2 | 1 | 64\|32 |

**Table S7:** Performance metrics for ordinal regression models predicting withdrawal severity.

| Model | RPSS | RPSS lower 2.5 | RPSS upper 97.5 | overall F1 | overall accuracy |
| --- | --- | --- | --- | --- | --- |
| All mice | 0.6244 | 0.4873 | 0.7475 | 0.79 | 0.8 |
| 129S1/SvImJ | 0.3408 | 0.0341 | 0.6011 | 0.56 | 0.64 |
| BALB/cJ | 0.3452 | -0.133 | 0.7155 | 0.63 | 0.62 |
| C57BL/6J | 0.7621 | 0.6125 | 0.8825 | 0.81 | 0.83 |
| DBA/2J | 0.551 | 0.1494 | 0.8871 | 0.73 | 0.82 |

## Notes

https://github.com/jacobbeierle/OW_analysis

### Competing Interest Statement

The authors have declared no competing interest.

